# Assembly-active and -inactive forms of HBV capsid protein provide distinctly different binding sites for capsid assembly modulators

**DOI:** 10.64898/2026.05.13.724798

**Authors:** Liam W. Scott, Carolina Pérez-Segura, Jodi A. Hadden-Perilla, Adam Zlotnick

## Abstract

In an infection, Hepatitis B Virus (HBV) core protein (HBc) normally assembles into icosahedral capsids. Capsid Assembly Modulators (CAMs) are direct acting antivirals that induce HBc mis-assembly and are the subject of active research and development. Two versions of HBc are used in structural studies of CAM-HBc complexes: Cp150 and Cp149-Y132A. Cp150 forms empty icosahedral capsids that are structurally indistinguishable from those found in virions. The Y132A mutation of Cp149 leads to an assembly defective soluble protein that crystalizes as flat hexagonal sheets, where the hexagons resemble icosahedral quasi-sixfold vertices. In this study, we compare structures of CAM-bound Cp150 to CAM-bound Cp149-Y132A. In capsids, the residues forming the CAM site shift to match the structure of bound CAMs, an induced fit. In Cp149-Y132A crystals, CAM sites show little structural adjustment in response to different CAMs binding. In turn, the array of residues that interact with CAMs varies from CAM to CAM in capsid structures but remains nearly constant in Cp149-Y132A crystals. These results illustrate important differences between CAM binding in Cp149-Y132A and Cp150 structures that will contribute to future CAM design.

## Introduction

Hepatitis B virus (HBV) chronically infects as many as 300 million people and results in ~1.1 million deaths per year [1]. Highly effective preventative vaccines are available, however, these vaccines have no effect on chronic HBV infection [2]. A functional cure of chronic HBV infection is defined as undetectable levels of HBV surface antigen (HBsAg) and viral DNA in patient sera after a finite treatment period [3-5]. Current standard of care for patients with chronic HBV infection is treatment with nucleoside analogs or pegylated interferon-alpha [6, 7]. These treatments reduce HBV replication and patient symptoms. However, they have very low rates of functional cure; viral recurrence is common upon end of treatment. Thus, new treatments are needed to achieve functional cure of chronic HBV infection [3-5].

The core of an HBV virion has a ~36 nm diameter capsid comprised of 120 homodimers of HBV core protein (HBc) arranged with T=4 icosahedral symmetry (**Figure 1**) [8]. In a T=4 capsid, HBc adopts four different quasi-equivalent conformations, chains A, B, C, and D. HBc dimers are labeled as AB or CD dimers in capsid structures (**Figure 1a-c**) [9-11]. During capsid assembly *in vivo*, 120 HBc dimers assemble around reverse transcriptase (RT) bound HBV ssRNA; notably, only ~10% of capsids assembled *in vivo* are filled with ssRNA while the remainder are empty [12]. We suggest that the basis for this division is that capsid assembly can be nucleated by the RNA-RT complex or spontaneously by HBc. After capsid assembly, this ssRNA is reverse transcribed inside the capsid into relaxed circular partially double stranded DNA (rcDNA). This complex is the infectious core, which has two possible fates: The core can be delivered to the host nucleus to maintain infection, where host machinery repairs the released HBV genome into covalently closed circular DNA (cccDNA). This cccDNA minichromosome acts as the source of HBV RNAs within chronically infected cells [5, 7, 13, 14]. Alternatively, cores can be surrounded by a layer of lipid and surface protein (HBsAg) to form infectious HBV virions. These virions are secreted from the host cell, where they go on to infect new cells [15-17].

**Figure 1:**
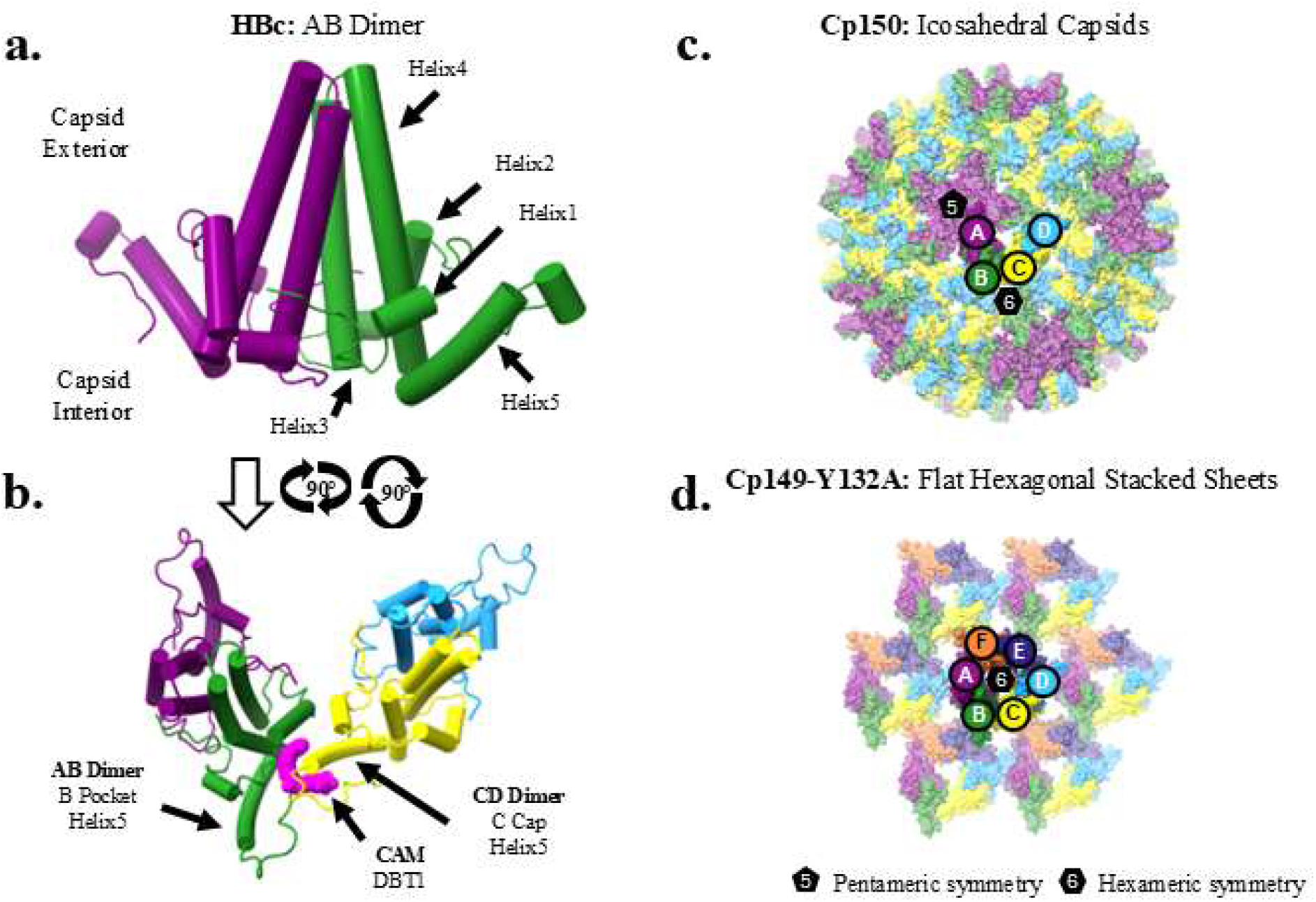
Quaternary structures of HBc and the CAM binding site. (**a**) A Cp150 AB homodimer, residues 1-141, from a T=4 capsid (PDB:6WFS). (**b**) Top view of HBc dimer of dimers. This is the icosahedral asymmetric unit. The CAM site is occupied by DBT1 (pink). Quaternary superstructures of HBc used to study CAM/HBc binding: (**c**) 120-dimer T=4 Cp150 capsid (PDB:6WFS), and (**d**) a sheet from a Cp149-Y132A crystal (PDB:5WRE).

Full length HBc (Cp183) has an assembly domain from residues 1-149, and an unstructured nucleic acid binding domain from 150-183. The nucleic acid binding domain of Cp183 is not necessary for the assembly capabilities of HBc and can be removed, resulting in the Cp149 construct. Cp149 can be recombinantly expressed and purified from *E. coli* and then assembled into empty HBc capsids [8, 18, 19]. In the assembly domain, helix3 and helix4 form the intra-dimer interface, creating a four-helix bundle spike that dots the surface of the HBc capsid (**Figure 1a-c**). The inter-dimer interaction is dominated by helix5. In one subunit the N-terminal half of helix5 forms a groove with helix2. The C-terminal half of helix5 from the neighboring subunit fills that groove with Y132-V124, which complements a notch at the end of the groove (**Figure 1b, Figure 2a-b**). Thus, helix5 of an HBc monomer acts as the “pocket” of one dimer and the “cap” for its neighboring dimer. This pocket-cap interaction is critical for proper geometric organization of HBc capsids.

**Figure 2:**
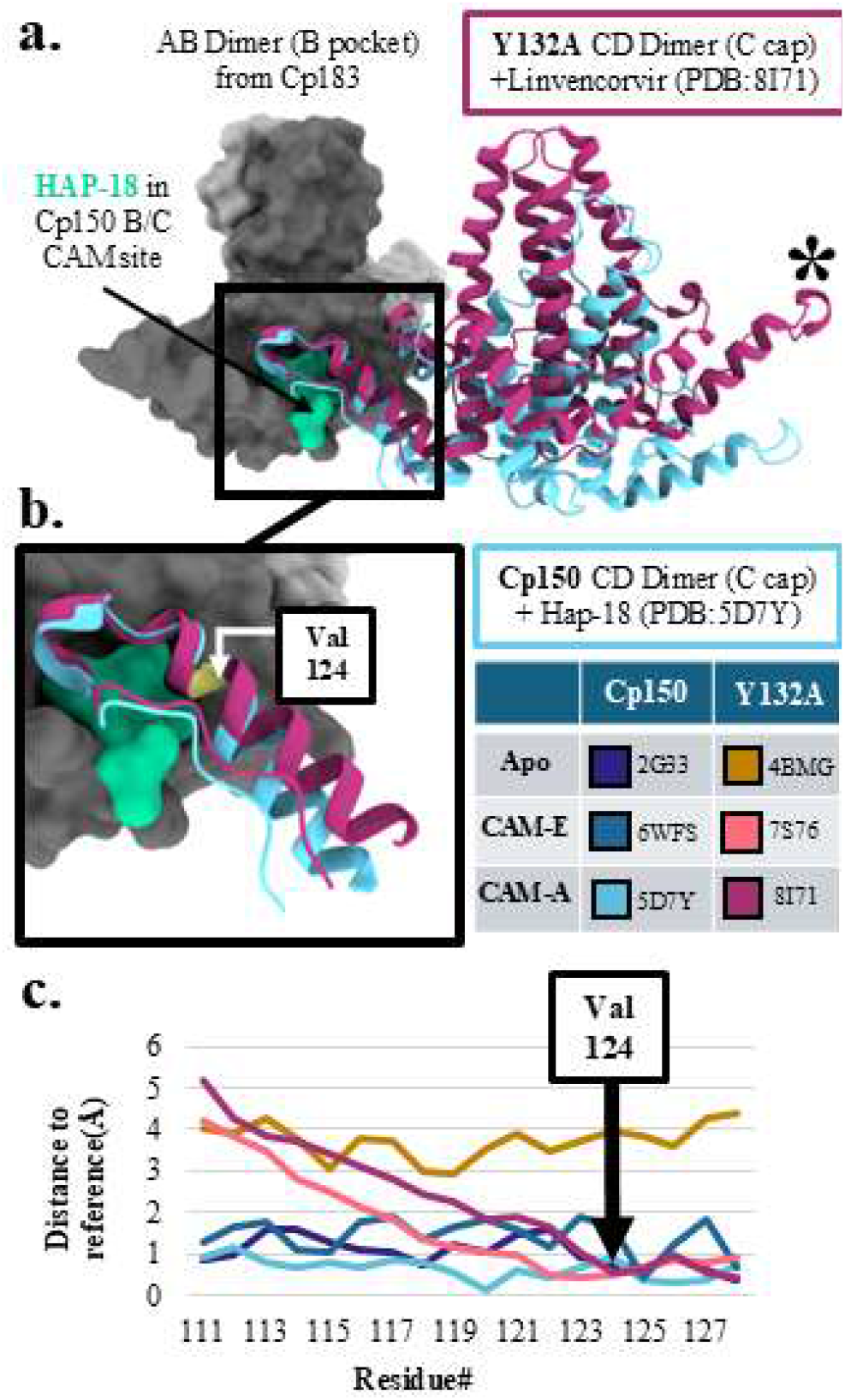
The geometry of the CAM pocket capping dimers is different for Cp149-Y132A and Cp150. (**a**) B pockets for (i) CAM-A bound Cp149-Y132A (PDBID: 8I71) and (ii) CAM-A bound Cp150 (PDBID: 5D7Y) aligned to apo Cp183 (PDBID:7OD4). Cp183 pocket dimer (A/B) displayed as a gray surface. Capping dimers are in distinctly different positions*. (**b**) Helix5 of the capping subunits for Cp150 and Cp149-Y132A overlap between Y132 and V124, which fills the groove above the CAM site. The two helices diverge at V124. (**c**) Distance of Cα of Cp150 and Cp149-Y132A structures from the Cp183 structure at capping helix 5. These data highlight differences that are structure dependent and CAM chemotype independent (**Supplemental Table 1**).

There are two assembly domain constructs that have made important contributions to structural studies of CAMs bound to HBc: Cp150 and Cp149-Y132A. Cp150 is a variant of Cp149 that has had its three native cysteines replaced with alanine and an additional cysteine appended to its C-terminus [20]. Cp150 assembles into empty, disulfide crosslinked capsids that retain the quasi-equivalent structure of wild-type capsids [21]. In contrast to Cp150, Cp149-Y132A is a single amino-acid mutation of Cp149 that significantly reduces the hydrophobic surface area buried at inter-dimer interfaces. This makes Cp149-Y132A highly resistant to assembling into capsids. However, Cp149-Y132A can be crystalized as stacks of flat hexagonal sheets, where the hexagons resemble icosahedral quasi-sixfolds (**Figure 1d**) [22, 23]. In keeping with their hexagonal organization, Cp149-Y132A chains in the crystallographic asymmetric unit are named A, B, C, D, E, and F.

Capsid assembly modulators (CAMs) are a promising class of drug towards achieving functional cure of chronic HBV infection (see **Supplemental Table 1**) [20, 24-42]. CAMs bind at the inter-dimer interface of HBc, between the pocket of one chain and the cap of the neighboring chain (**Figure 1b, Figure 2a-b**) [20]. Herein, we shall refer to a specific CAM site by its pocket/cap name, e.g. the CAM site between the B chain pocket and the C chain cap is “B/C” (**Figure 1b**). CAMs act by strengthening the association energy between dimers and accelerating assembly of HBc. There are two major classes of CAM: CAM-Aberrant (CAM-A) molecules result in non-capsid, dysfunctional HBc structures. In contrast, CAM-Empty (CAM-E) molecules generate minimally distorted, empty capsids [43]. In both cases, CAMs appear to facilitate spontaneous nucleation of assembly in lieu of nucleation by RNA-RT, leading to reduced secretion of virions in culture and *in vivo* [24, 30, 44]. Some CAMs have an additional mechanism of action where they disrupt mature DNA filled virions, preventing infection of new cells [36]. This shows that CAMs can attack HBc at multiple stages of the HBV life cycle. CAM-As are of particular interest for functional cure development; CAM-A treatment of HBV infected mice and cell cultures have been shown to trigger apoptosis of infected cells [24]. Such apoptosis is a potential route for removing HBV nucleic acid from the host during chronic HBV infection. However, this effect has only been observed when CAM-As are applied during a highly active infection. Thus, further development of CAM efficacy is required to broaden their treatment window for functional cure of chronic HBV infection [44].

Since CAMs bind at the interaction interface between HBc dimers, the angle of interaction between HBc dimers is critical for defining the shape of the CAM binding site. However, Cp149-Y132A’s flat sheets and Cp150’s spherical capsids require different angles of interaction between subunits to form (**Figure 1c-d, Supplemental Figure 1**). Furthermore, there are four quasi-equivalent folds of HBc in capsid structures, which result in four different quasi-equivalent CAM sites: A/A, B/C, C/D, and D/B [35]. In contrast, the hexagonal repeat of Cp149-Y132A in its crystallographic unit cell results in six quasi-equivalent CAM binding sites: A/D, B/C, C/F, D/E, E/B, F/A (**Methods**) [41]. These observations together suggest that there are critical differences in the CAM binding sites of Cp150 and Cp149-Y132A structures. In this study, we performed comparative structural analysis of available structures of apo and CAM-bound Cp150 to apo and CAM-bound Cp149-Y132A (**Supplemental Table 1**). We observe critical differences between CAM-induced conformational changes in Cp149-Y132A and Cp150 structures. These results are of importance for interpreting binding activity and guiding structure-based design of new CAMs.

## Results

### Dimer-Dimer Angle of Interaction

To provide a common perspective, we chose a high-resolution RNA-filled capsid as a reference structure, PDB:7OD4 [10]. To the B pocket of this reference structure, we aligned the B pockets from apo and CAM-bound B/C CAM sites of Cp150 and Cp149-Y132A structures (**Figure 2a, Supplemental Table 1**). Specifically, we describe the pocket as selected residues from the pocket chain of the CAM site: helix2 through helix3 (23-37), and the C-terminal half of helix4 through the N-terminal half of helix5 (100-120). These pocket residues were chosen because they show minimal conformational variation from structure to structure (**Supplemental Figure 2**). The alignment of these selected residues from the B pocket of the Cp183 reference to B pockets from five CAM-bound Cp150 capsids and ten CAM-bound Cp149-Y132A sheets (**Supplemental Table 1**) have RMSDs that vary from 0.356-0.758 Å. For comparison without bound CAMs, the B pocket of apo Cp150 (PDB:2G33) has an RMSD of 0.547 Å between its quasi-equivalent pockets; apo Cp149 (PDB:1QGT) B pockets have an RMSD of 0.326 Å; apo Cp149-Y132A (PDB:4BMG) pockets have an RMSD of 0.365 Å. Of note, CAM-bound Cp149-Y132A structures only begin to diverge from capsid structures as one looks further from the pocket chain residues, e.g. approaching the C-terminal half of helix 5, which is where the capping region of the neighboring CAM site begins (**Supplemental Figure 2**). These data emphasize that the aligned *pocket* component of the CAM site is highly similar across different HBc structures.

In our alignment of the structurally conserved pocket dimers, the capping dimers adopt different relative orientations (**Figure 2**). CAM-bound Cp149-Y132A capping dimers and their pocket dimers create a flat plane, constrained by crystal packing. Surprisingly, a different position is seen in the apo Cp149-Y132A structure. In contrast to CAM-bound Cp149-Y132A capping dimers, the capping subunits from apo Cp150 and CAM-bound Cp150 are shifted downwards in relation to this plane; the offset interaction is expected in order to conform to the curvature of a capsid structure. This is displayed with representative structures in **Figure 2a**, which highlights the difference in capping subunit orientations between Cp149-Y132A and Cp150 structures that are bound with similar CAMs. These differences are most easily visualized at the tip of the capping dimer that is distal from the CAM site (the asterisk in **Figure 2a**).

The relationship between capping and pocket subunits shows differences in the inter-dimer interactions between Cp149-Y132A sheet and Cp150 capsid structures, both with and without bound CAMs (**Figure 2a-b**). To demonstrate these interactions, we calculated distances between C-alphas in capping helix5 of the Cp183 reference structure and equivalent residues in the aligned CAM bound HBc structures (**Figure 2c**). Except for apo Cp149-Y132A, all tested structures are closely aligned near the CAM site. However, distal from the CAM site, CAM-bound Cp149-Y132A structures diverge from CAM bound capsid structures. The fulcrum of this divergence is near residue V124 (**Figure 2b-c**). We also observe in this alignment that the entire apo Cp149-Y132A structure is shifted by several Ångstroms compared to other structures in this study, indicating a distinct quaternary interaction.

To quantify the differences in CAM site geometry, we calculated the angle-of-interaction (AOI) between capping and pocket subunits (**Figure 3**). We define this AOI as the angle between helix 5 (residues 111-127) of capping and pocket chains (**Figure 3a-b, Methods eq. 1**). This AOI between dimers is fundamental to describing the shape of the CAM site. For the ten CAM-bound Cp149-Y132A structures, the six quasi-equivalent CAM sites have a similar AOI of 42.6° ± 1.3° (Ave. +/-st. dev.). For the five CAM-bound Cp150 capsid structures, the AOI between chains for A/A CAM sites is 42.0° ± 1.0° and C/D CAM site is 41.8°±1.3°. However, the AOI for the B/C CAM site is 34.4° ± 2.6° and D/B CAM sites are 37.5°±1.4° (**Figure 3c**). This indicates systematic differences in the architecture of Cp150 and Cp149-Y132A sites.

**Figure 3:**
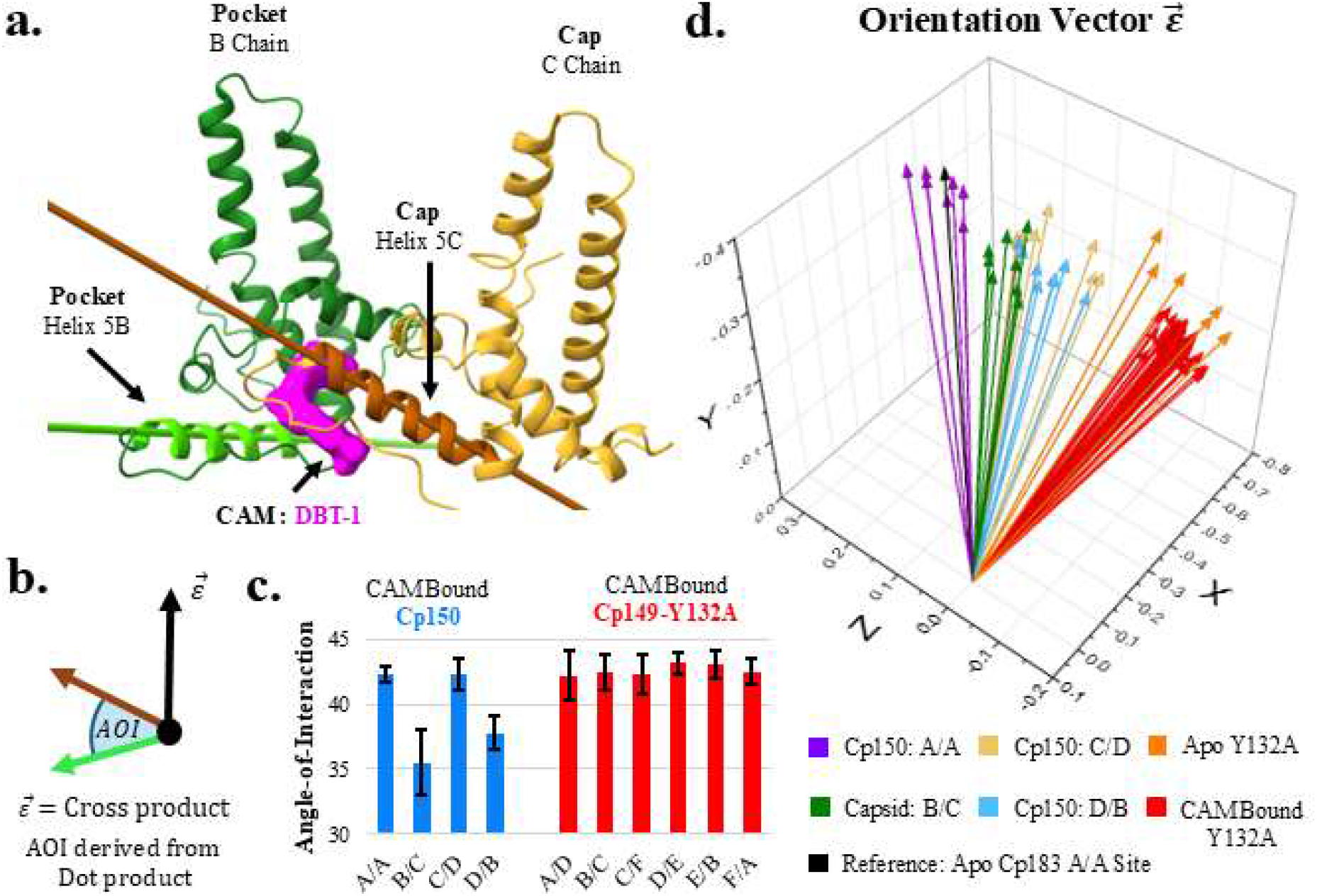
Dimer-dimer geometry for Cp150 and Y132A. (**a**) Vectors for the pocket helix 5 (lime green) and the capping helix 5 (brown). (**b**) Schematic of vector calculations: The angle between pocket and cap vectors is the angle-of-interaction (AOI). The cross product of the helix vectors 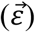 reports the orientation of the CAM site in relation to the aligned capsid A/A site. (**c**) Average dot products of all CAMs for quasi-equivalent sites. Error bars are one standard deviation. N=5 for Cp150 capsids and N=10 for Cp149-Y132A crystals. (**d**) Epsilon vectors where all pockets are aligned with the apo-Cp183 capsid A/A site. (**Supplemental Table 1**)

We also compared CAM site orientations with respect to the capsid. MD simulations have shown that both CAM-As and CAM-Es can distort quasi-sixfolds at their edges [45]. Here we provide a local measure of this change: The CAM site orientation vector 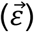. This vector is the cross-product of pocket helix5 and capping helix5 vectors (**Figure 3a-b, Methods eq. 2**). In this analysis, the pocket for each CAM-bound and apo structure was aligned with that of the reference using the same method as in **Figure 2**. We used the A/A site of the RNA-filled, CAM-free capsid (PDBID: 3J2V) for the reference for clarity of data presentation. Again, For Cp149-Y132A structures we observe a narrow cluster of 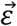, as might be expected for proteins constrained by crystal packing. Conversely, 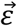 for Cp150 capsids are spread across a broad range that is separated from Cp149-Y132A structures based on the quasi-equivalence of the CAM site (**Figure 3d**).

To provide a more intuitive quantification of the differences in **Figure 3d**, we calculate the angle between 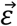 of a given sample and the A/A reference 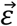 vector (**Methods eq. 1**) resulting in a “CAM site orientation angle” (**Figure 4a**). This angle describes the orientation of CAM sites relative to the A/A site of the capsid. All Cp149-Y132A site 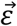 are closely clustered at ~38° from the reference, while Cp150 CAM site 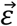 differ from the reference by anywhere from 2-20°. CAM-bound and apo Cp150 site 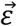 are found in three overlapping clusters, A/A, B/C, and C/D+D/B, illustrating the variability of quasi-equivalence. Again, apo Cp149-Y132A quasi-equivalent site 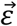 are further separated from both CAM bound Cp149-Y132A, CAM-bound Cp150, and apo Cp150 groupings (**Figure 4b-c**).

**Figure 4:**
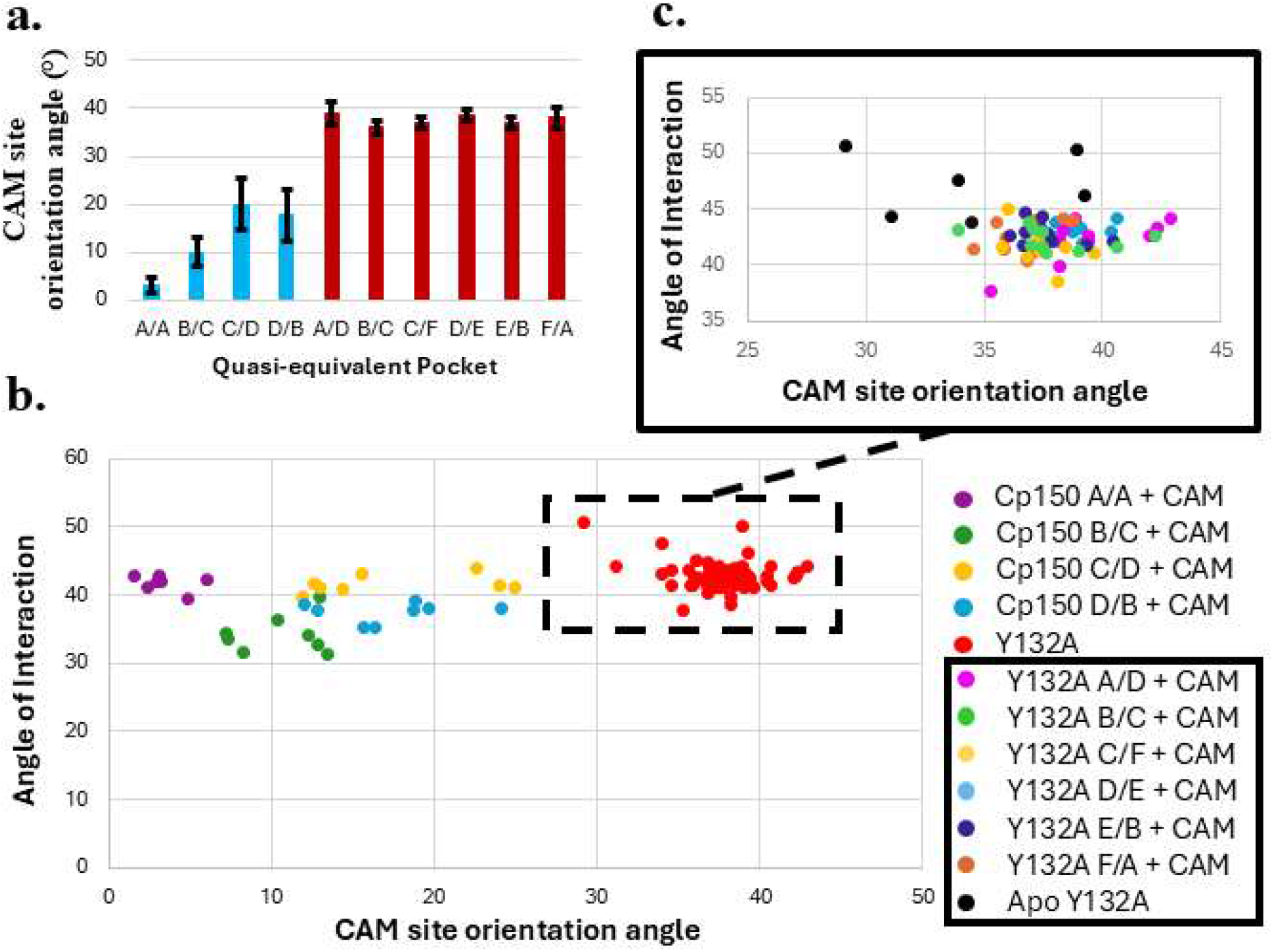
The CAM site orientation angle is sensitive to quasi-equivalence and CAM binding. (**a**) The CAM site orientation angle, graphed as the average of each quasi-equivalent site in measured structures. (**b**) Correlation of the AOI from **Figure 3c** to the CAM site orientation angle of **Figure 4a** colored by structure and quasi-equivalent pocket. (**c**) Inset shows zoom of only Cp149-Y132A sites. (**Supplemental Table 1**)

### CAM Site Topography

The shape and size of the CAM site are fundamental to describing CAM-HBc interactions (**Figure 5, Supplemental Table 1**). To calculate CAM site shape and size for comparison between Cp149-Y132A and Cp150 structures, we used *Measure volinterior* with fuzzy boundary conditions within Visual Molecular Dynamics (VMD) (**Methods**) [46, 47].

**Figure 5:**
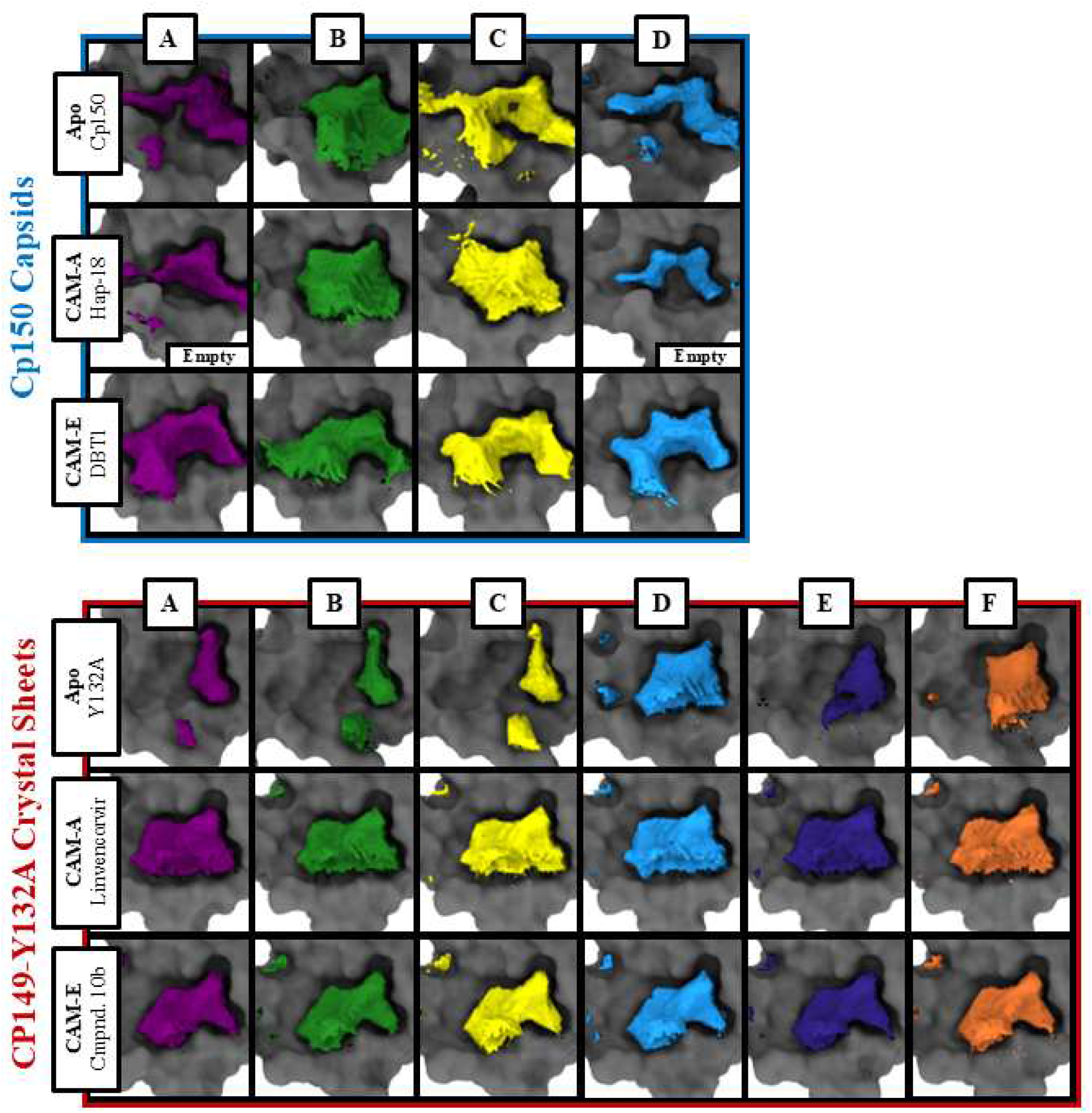
CAM site volumes. Quasi-equivalent CAM site shapes for Cp150 capsids (upper panel) and Cp149-Y132A (lower panel) overlaid on the corresponding pockets for apo, CAM-A (HAP-18,Linvencorvir), and CAM-E (DBT1, compound 10b) bound sites. The capping chains are hidden for clarity. Shapes were determined using *Measure volinterior* with fuzzy boundary conditions implemented in VMD.

Cp149-Y132A hexamer sheets and Cp150 capsids have different structural responses to CAM binding (**Figure 5**). In apo Cp150 capsids, the four quasi-equivalent sites have different shapes. When bound by CAMs, we observe that Cp150 CAM sites alter their shape, conforming to the structure of the bound CAMs. Bulky heteroaryldihydropyrimidines (HAPs) fill only B/C and C/D quasi-equivalent CAM sites in Cp150 structures, resulting in correspondingly large volumes. The unfilled A/A and D/B pockets become smaller in comparison to their apo counterparts. In contrast to HAPs, dibenzothiazepine1 (DBT1) binds to all quasi-equivalent pockets; the pockets become crescent shaped to conform with the shape of the bound DBT. This exemplifies how Cp150 CAM sites can adapt to the shape of the bound drug, an induced fit that is modulated by quasi-equivalence.

In the CAM sites of the apo Cp149-Y132A sheet structure (PDB:4BMG) we find the available volume adopts several different shapes across the six quasi-equivalent sites: three sites are small and similar to one another, while the other three sites are larger and vary in their shape. We note that the smaller Cp149-Y132A sites result from differences in the distances between Trp102 of the pocket subunit and Pro129 of the capping subunit: Trp102 and Pro129 are 4 Å apart in smaller apo Cp149-Y132A pockets, and up to 8 Å apart in larger Cp149-Y132A pockets. When filled by CAMs, Cp149-Y132A quasi-equivalent sites all adopt a similar, irregular shape (**Figure 5**). The chemical structures of the CAMs bound in available Cp149-Y132A crystal structures have a wide range of shapes. However, CAM structure has only a modest influence on the shape of CAM site volume in Cp149-Y132A structures, especially when compared to Cp150 structures (**Supplemental Figure 3**). Cp149-Y132A crystal structures are generated by first growing apo Cp149-Y132A crystals and then soaking CAMs into those crystals. Therefore, structural changes may be limited by crystal packing constraints.

Volumes for the CAM sites in capsids are sensitive to both CAM shape and quasi-equivalence (**Figure 6**). In apo Cp183 volumes for quasi-equivalent sites range from 85-270 Å^3^, apo Cp150 quasi-equivalent site volumes range from 200-360 Å^3^, and apo Cp149 site volumes range from 140-290 Å^3^. Apo Cp149-Y132A quasi-equivalent sites vary from 100-250 Å^3^. In CAM-bound Cp150 capsid structures, CAM site volumes range from 75-450 Å^3^ across filled and unfilled quasi-equivalent sites. However, in CAM bound Cp149-Y132A structures, site volumes are 225 ± 50 Å^3^, a much narrower range, and all sites are CAM filled. All Cp149-Y132A CAM structures show strong similarities in both CAM site shape and positioning. This contrasts with the substantial differences in pocket shape and size between CAM-bound Cp150 structures (**Supplemental Figure 3**).

**Figure 6:**
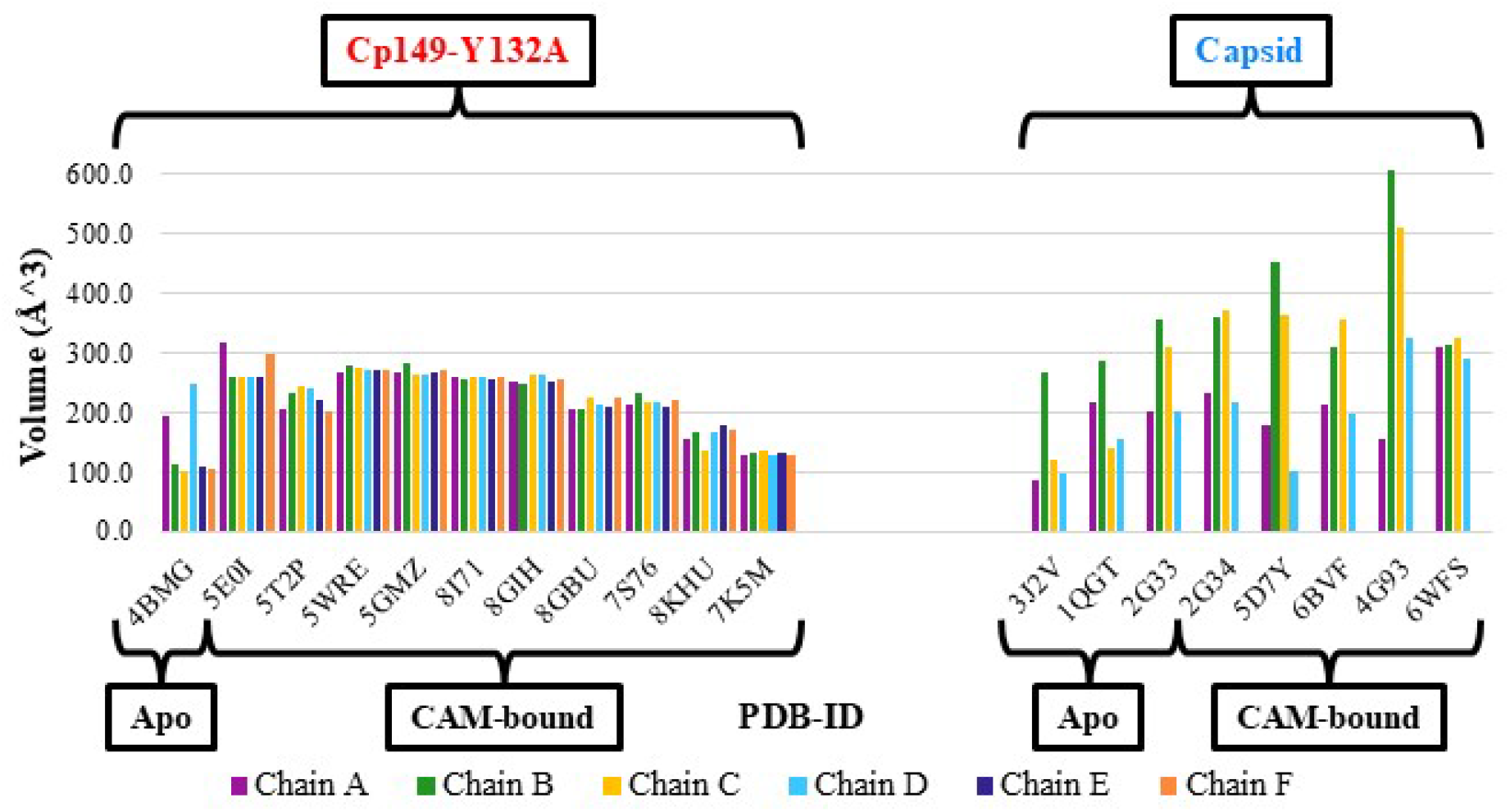
CAM site volumes. CAM site volumes for apo Cp183, apo/CAM-bound Cp150, apo Cp149 and apo/CAM-bound Cp149-Y132A structures. Volumes were calculated from shapes composed of voxels that have been determined to have ≥90% of rays cast from them occluded by protein surfaces, as determined by *Measure volinterior* with fuzzy boundary conditions in VMD. See **Supplemental Table 1** for further structure information.

The differences in CAM site shapes between Cp150 capsid and Cp149-Y132A sheet structures suggest that different amino acids are involved in Cp-CAM interactions. We identified residues that hydrogen bond or interact with bound CAMs in Cp149-Y132A and Cp150 structures (**Methods**). We find that Cp150 structures have diverse patterns of residues that interact with different CAM chemistries and quasi-equivalent sites (**Supplemental Table 2**). In contrast, Cp149-Y132A CAM interactions are nearly identical between different CAM sites and CAM chemistries (**Supplemental Table 3**). We identified eight amino acid residues that interact with every CAM in Cp149-Y132A CAM structures (**Figure 7**). In contrast, only one residue interacts with every CAM in CAM-bound Cp150 sites: Thr128 of the capping chain. Indeed, capping chain Thr128 is the *only* residue found to interact with CAMs at every quasi-equivalent site with every CAM in *both* Cp150 and Cp149-Y132A structures.

**Figure 7:**
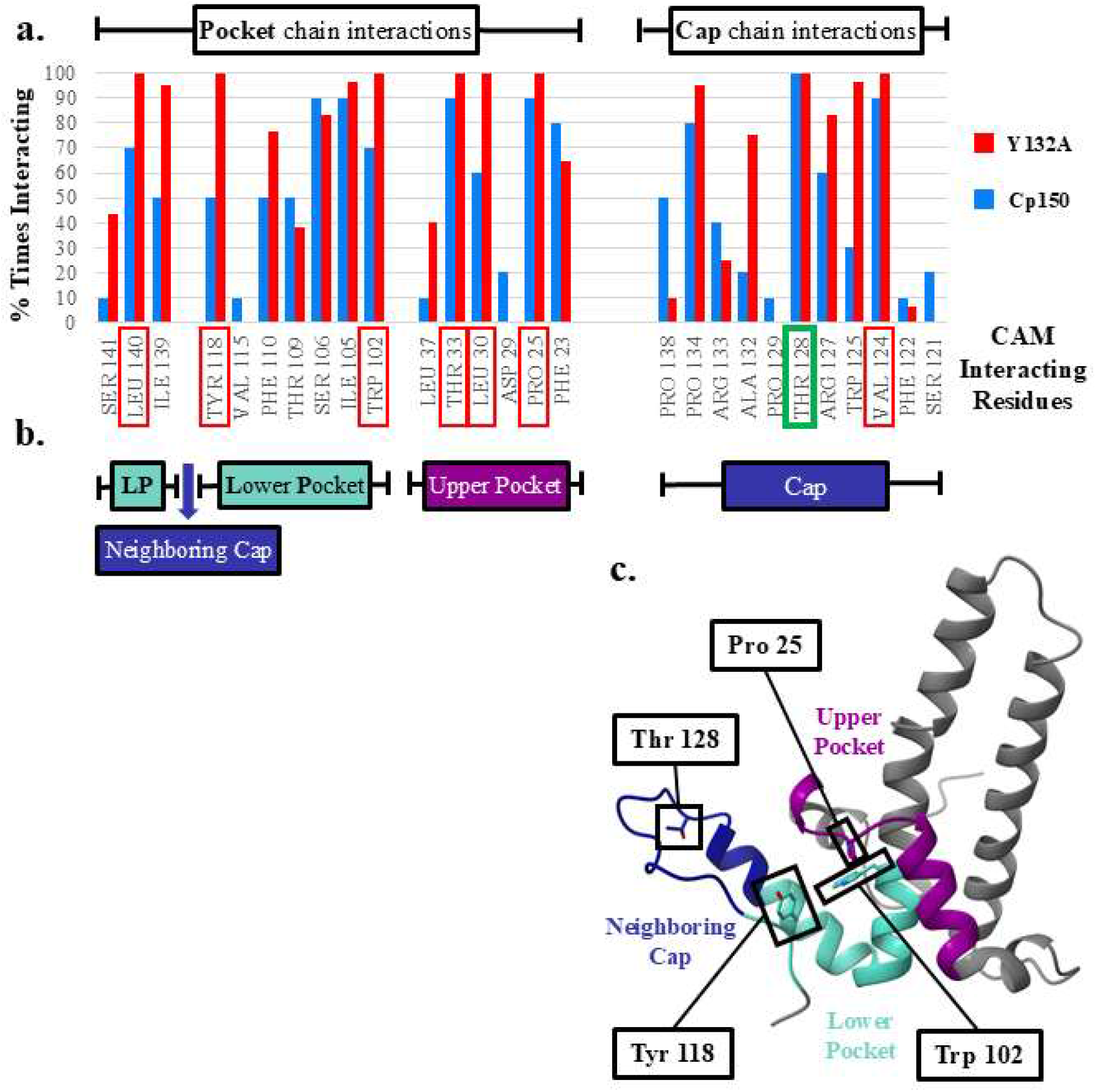
HBc residues that interact with bound CAMs. **(a)** HBc amino acid residues that interact with bound CAMs. Interactions represented as the percentage of times that residue interacts with a CAM across either ten Cp150 CAM bound quasi-equivalent sites or sixty CAM bound Cp149-Y132A quasi-equivalent sites. Residues that always interact in Cp149-Y132A models are highlighted in red. Thr128 (green) interacts with every CAM in every structure (**b**) Infographic correlating CAM interacting amino acids to regions of the HBc dimer-dimer interaction. (**c**) Structure of HBc colored according to **b**, with the most regularly interacting amino acids highlighted.

We also observe that Cp149-Y132A structures have a common hydrogen bonding pattern in their quasi-equivalent sites, while Cp150 structures have large variations in their hydrogen bonding patterns (**Supplemental Table 4-5**). For example, all Cp149-Y132A CAM-bound structures formed hydrogen bonds with the pocket residue Trp102 (60 CAM bound Cp149-Y132A sites total). In contrast, Trp102 formed a hydrogen bond with only one HAP variant in one quasi-equivalent pocket in Cp150 (out of 20 CAM bound Cp150 sites total). We note that the resolution of capsid structures overall is lower than that of Cp149-Y132A crystal structures, which may contribute to some of the differences in interacting residues seen between Cp150 and Cp149-Y132A. However, prior measurements in this study support our observation that Cp150 structures have variable interactions with different bound CAMs, while Cp149-Y132A structures have highly similar interactions with different bound CAMs.

## Discussion

Cp149-Y132A structures have different CAM-HBc interactions than those observed in Cp150 capsid structures. When a CAM binds to a site in a Cp150 capsid, the amino acids residues adjust to complement the structure of the CAM. Recent results suggest that CAM binding in one quasi-equivalent Cp150 CAM site can cause shifts in the shape of neighboring quasi-equivalent CAM sites [48]. This effect can yield binding with negative co-operativity: Filling one quasi-equivalent CAM site with a CAM causes shifts in other nearby sites that makes them less favorable for CAM binding [45, 48]. We note that in the case of molecular dynamics simulations of CAM bound capsids, positive cooperativity can also be observed [36, 45]. Further, HDX-mass spectrometry experiments confirm that CAM binding can significantly change the dynamics of Cp150 structures [49]. These studies, in conjunction with our own, demonstrate that binding a CAM induces structural changes that propagate across a capsid. In contrast to Cp150, Cp149-Y132A structures exhibit a single response to CAM binding, regardless of CAM chemistry and structure. We propose that this limited response may be an artifact of crystal contacts restraining the mobility of HBc in response to CAM binding.

Biochemically, Cp150 structures have a wide variety of pocket shapes and sets of residues that interact with different bound CAMs. In contrast, CAM-bound Cp149-Y132A CAM sites and their CAM-interacting residues are highly similar to each other. Of note, many CAM resistance mutations identified in HBc fall within regions 23-37 (Helix2) and 106-128 (from the base to the middle of helix5) [50-52]. This observation is consistent with CAM-bound Cp150 and Cp149-Y132A structures, where these regions nearly always interact with CAMs. The CAM site orientation angle (**Figure 4**) reflects these differences, with Cp150 CAM sites separated by quasi-equivalence while CAM-bound Cp149-Y132A structures cluster together away from Cp150 CAM sites. We suggest that CAM site orientation angle will be a useful index for relating CAM binding to structural disruption in future studies.

In capsids we found different patterns of hydrogen bonding residues dependent upon quasi-equivalence and CAM chemistry. In Cp149-Y132A crystals, CAM hydrogen bonding patterns are similar across quasi-equivalent chains and CAM chemistries but different from those in Cp150. The most striking example of this is Trp102, which in Cp149-Y132A forms hydrogen bonds with every CAM type in every Cp149-Y132A quasi-equivalent CAM site in tested structures. In comparison, this hydrogen bond is rarely seen in Cp150 capsids (**Supplemental Table 4-5**). We also note that the smaller CAM sites available in apo Cp149-Y132A are a result of interactions with Trp102. This further highlights Trp102 as having different interactions in Cp150 and Cp149-Y132A structures.

CAMs are optimized for their ability to induce capsid assembly and to disrupt normal HBc dimer-dimer interactions. Cp149-Y132A crystal structures loosely resemble misassembled Cp protein from CAM-A treatment that are enriched for hexamers [36, 53, 54]. However, the Y132A mutation changes the geometry of protein-protein interactions. Recent structural and molecular dynamics studies further support the conclusion that Cp149-Y132A structures have alternate protein-protein interactions which modify the shape and accessibility of the CAM site in comparison to capsids [55]. In contrast, Cp150 capsids appear to be accurate models for HBc capsids [21, 56]. Overall, these studies show that Cp150 and Cp149-Y132A structures provide different views of CAM-HBc binding interactions. Thus, we suggest that future structure-guided design of CAMs should consider the inherent differences between capsid and Cp149-Y132A structures.

## Methods

PDB files were analyzed using ChimeraX [57-59]. Since the last modeled residue for these structures varies, all residues beyond 141 were removed from the structures for consistency across structures. Hydrogens were then added through ChimeraX’s addH function prior to further structural analysis, which calculates histidine protonation utilizing local environment. All other amino acids are protonated by addH based on standard physiological conditions.

To compare CAM sites in Cp149-Y132A sheets and Cp150 capsids, we initially chose the B/C CAM binding site of an RNA-filled Cp183 capsid as a reference, PDBID: 7OD4. This is a cryo-EM structure of an apo capsid that is lacking bound CAM. The RNA acts as a stabilizing agent and is a good model of a naturally occurring nucleic acid filled capsid. For capsids, chain naming is based on the standard definition of quasi-equivalence for a T=4 virus capsid. For Cp149-Y132A crystal structures, the labeling of subunits is arbitrary but is consistent across structures analyzed in this study. The exception to this is apo Cp149-Y132A, [22] in which the chain capping the B pocket is named “E” rather than “C”. Therefore, in apo Cp149-Y132A (PDB:4BMG) the names of the E and C chains was swapped, and the names of the D and F chain was swapped in order to line up apo and CAM-bound Cp149-Y132A structure nomenclature. As the repeats of the CAM site in the asymmetric unit of Cp149-Y132A crystals are not identical, Cp149-Y132A CAM sites are also described as quasi-equivalent. A list of the structures used for the following studies can be found in **Supplemental Table 1**. Examples of relevant CAM structures for various studies are displayed in **Supplemental Figure 3**.

For initial alignment of pockets, ChimeraX’s matchmaker function was used to align residues 23-37 and 100-120 of the pocket chain of the structures in question to the pocket chain of the reference structure. RMSD’s are reported by ChimeraX’s MatchMaker function. C-alpha distances reported in **Figure 2c** and **Supplemental Figure 1b** were calculated from the distance between C-alpha’s in the reference structure and the C-alphas of the structure in question.

To extract helix5 vectors for analysis, residues 111-127 were selected, and a vector was generated centered in the alpha carbons of those residues using ChimeraX’s higher-order structure axes tool. The angle between the pocket and capping helix5 vectors, the angle of interaction (AOI), was calculated from the dot product of the two vectors (**Equation 1**). The vector perpendicular to the plane formed by the pocket and capping vectors, the orientation vector 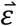, was calculated from their cross product (**Equation 2)**. These metrics are illustrated in **Figure 3b**.

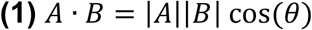

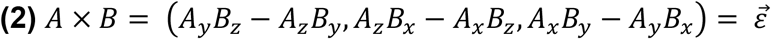

To determine CAM site shape and volume, the GPU implementation of *measure volinterior* with fuzzy boundary conditions, part of Visual Molecular Dynamics (VMD), was used [46, 47]. This function uses a ray-casting method to classify the space associated with ligand binding pockets based on the surrounding protein surface. Voxels of space are selected for pocket visualization based on the number of rays cast from them that strike the protein surface versus those that are not. This approach allows for generation of space filling models of protein pockets at various levels of confidence [47]. CAM site shapes visualized and volume values reported (**Figure 5-6**) using voxels that have 90% of rays cast from them occluded by the protein surface. It is important to note that selecting the appropriate protein surface parameters for your protein of interest in these calculations is critical. The protein surfaces for *measure volinterior* were generated based on CAM site residues (19-37, 98-102, 104-106, 108-111 of the pocket chain and 18, 120-138 of the capping chain) using VMD’s default Van der Waals radii and QuickSurf parameters of: RadiusScale = 1.2, Isovalue = 0.8, and GridSpacing = 0.5. The rays cast from each voxel was: Nrays = 64.

To analyze residues that interact with CAMs in HBc structures, ChimeraX’s “contacts” and “H-bonds” functions were used. To study contacts, all residues with a Van der Waals overlap 0.5 Å or greater with the bound CAM in question are listed as interacting residues (**Supplemental Table 4-5**). For H-bonds, a distance tolerance of 0.450 Å and an angle tolerance of 20° was used when identifying H-bonds between HBc and bound CAMs.

## Acknowledgements

AZ acknowledges funding from NIH R01-AI118933. JAH-P acknowledges funding from NIH award P20-GM104316-10.

## Supplemental Data

**Supplemental Table 1:**
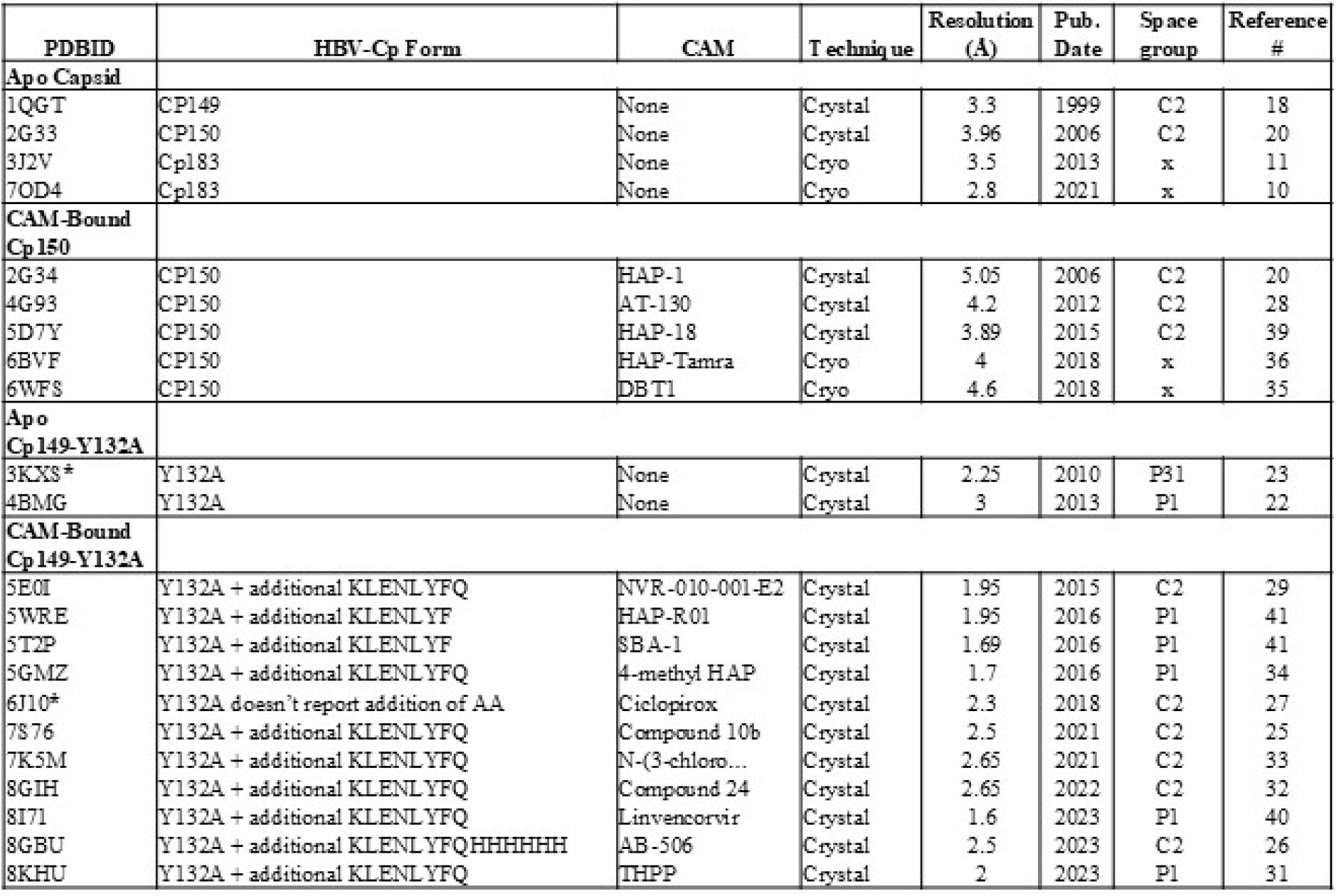
List of HBV-Cp structures used in study. 2G33 is used as “apo Cp150” in all studies and 4BMG is used as “apo Cp149-Y132A” in all studies. The only other CAM bound Cp149-Y132A structure that we found was 8W9G. 8W9G does not have an associated publication as of this time, and so it was not included in this study. 7OD4 was used as the alignment structure for **Figure 2** and **Supplemental Figure 2**. 3J2V A/A pocket was used to align data for **Figure 3-4**. 1QGT was used to align pockets for **Supplemental Figure 3**. 3J2V was utilized as an apo form of Cp183 to act as a reference to compare against in **Figure 6, Supplemental Figure 1**, and **Supplemental Figure 3**. * Structures not used in this study: 3KXS crystallized with a different packing arrangement from 4BMG and the other CAM bound Cp149-Y132A structures. 6J10 has a different chain naming nomenclature from all other Cp149-Y132A structures.

**Supplemental Figure 1:**
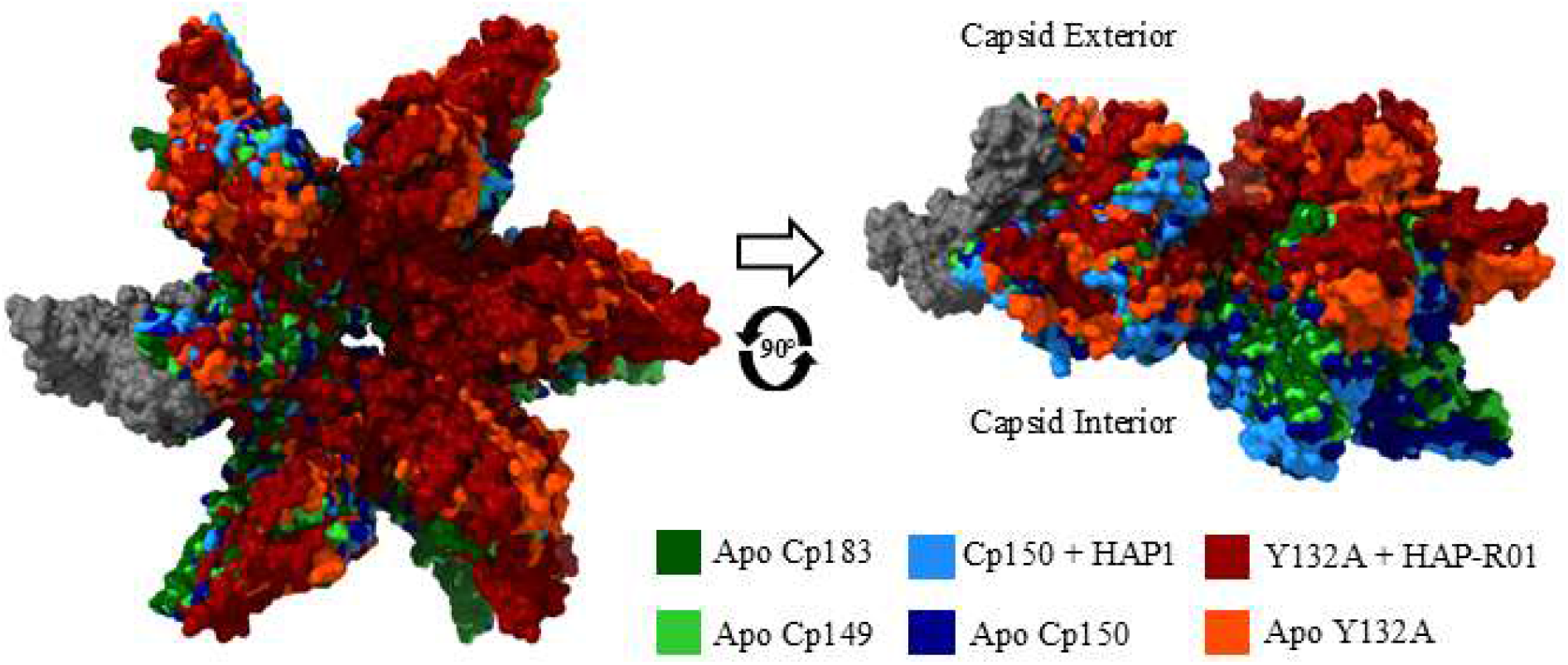
Hexameric units from capsids and Cp149-Y132A sheets have different curvatures. External/Top view (left) and side view (right) of an alignment of apo and CAM bound hexameric units from apo Cp183 (PDB:3J2V), apo Cp149 (PDB:1QGT), Cp150 (apo PDB:2G33, + CAM PDB:2G34), and Cp149-Y132A (apo PDB:4BMG, + CAM PDB:5WRE). Only the D chain (gray) was aligned using ChimeraX’s Matchmaker function. This alignment shows that hexamers of capsids have greater curvature than hexamers from Cp149-Y132A sheets, independent of CAM binding.

**Supplemental Figure 2:**
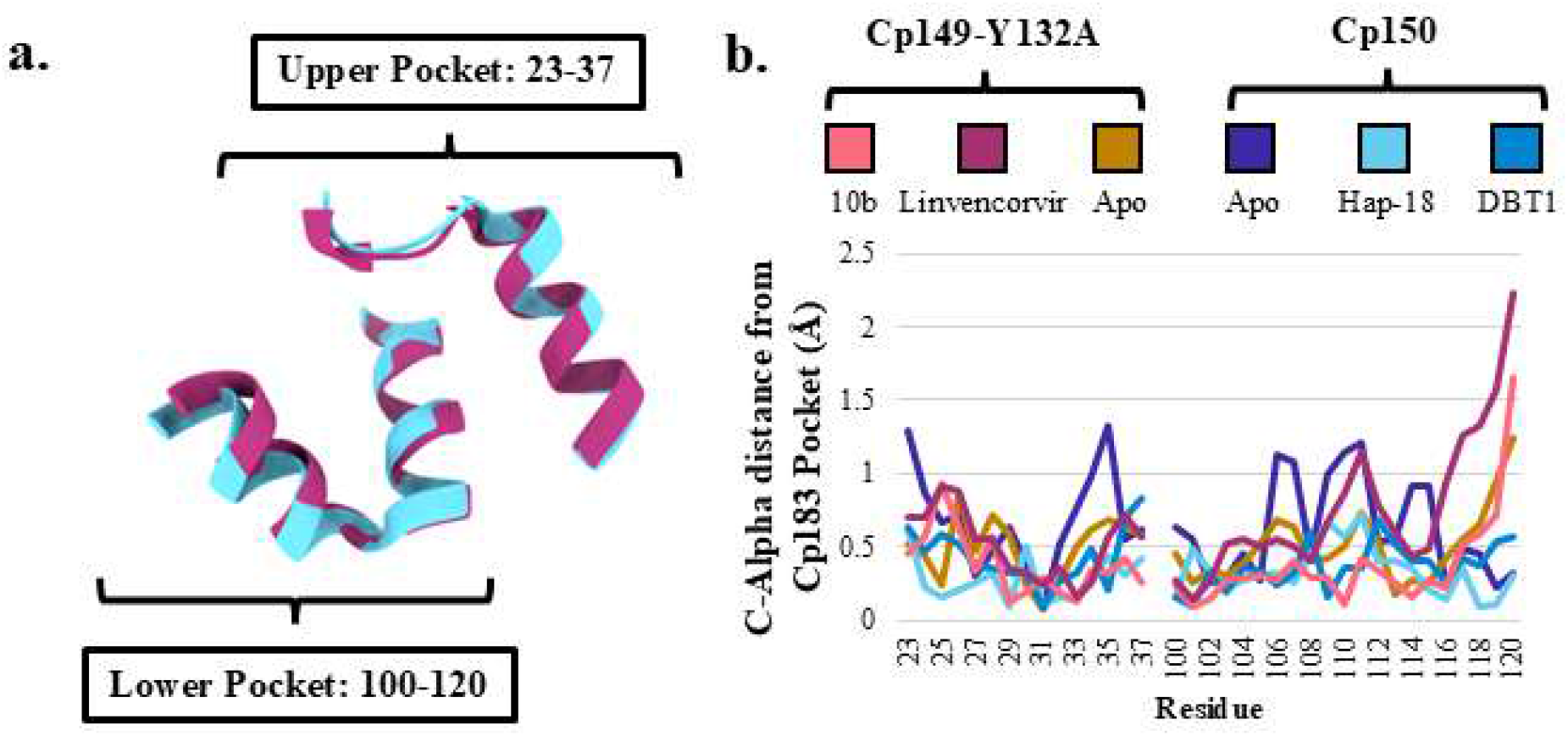
Aligned pocket alpha-carbons are highly similar across apo and CAM bound structures of Cp150 and Cp149-Y132A. (**a**) Alignment of residues 100-120 and 23-37 of Linvencorvir bound Cp149-Y132A (PDBID:8I71) and Hap-18 bound Cp150 (PDBID:5D7Y) to the B pocket of RNA filled, apo Cp183 (not shown, PDBID: 7OD4). (**b**) C-alpha to C-alpha distance of the aligned B pockets of several CAM bound Cp149-Y132A and Cp150 structures from the aligned-to apo Cp183 structure. Overall, samples do not vary more than 1.0 Å from the aligned-to pocket until approaching the portion of helix5 that acts as the cap for the neighboring pocket. (**Supplemental Table 1**)

**Supplemental Figure 3:**
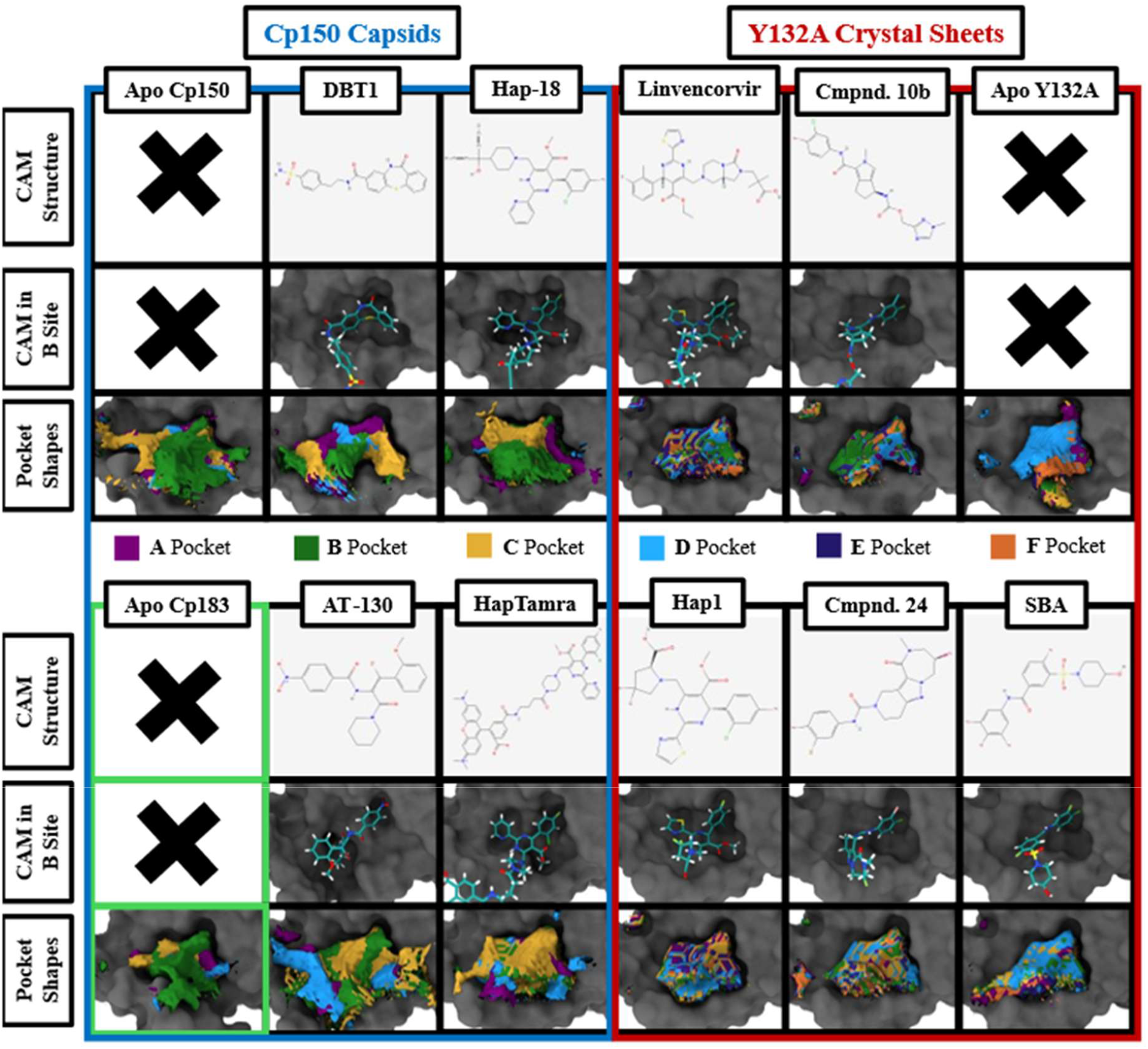
Aligned CAM site shapes are similar in Cp149-Y132A structures and variable in capsid structures. Prior to calculating CAM site shape and volume using VMD’s *Measure volinterior* with fuzzy boundary conditions, all structures were aligned to the B pocket of Cp149 (PDBID:1QGT) using residues 100-120 and 23-37. This allows for overlays of the CAM sites from different quasi-equivalent sites. Stick structures of a CAM displayed at top, an image of that CAM bound in the B site of its Cp150 or Cp149-Y132A structure with the B pocket displayed as a gray surface in the middle, and that same B pocket with shown with an overlay of all VMD calculated quasi-equivalent site shapes from that structure. We note that CAM site volume shapes are nearly identical across overlaid Cp149-Y132A quasi-equivalent sites, while Cp150 site volume shapes show variability in shape and orientation between quasi-equivalent sites. (**Supplemental Table 1**)

**Supplemental Table 2:**
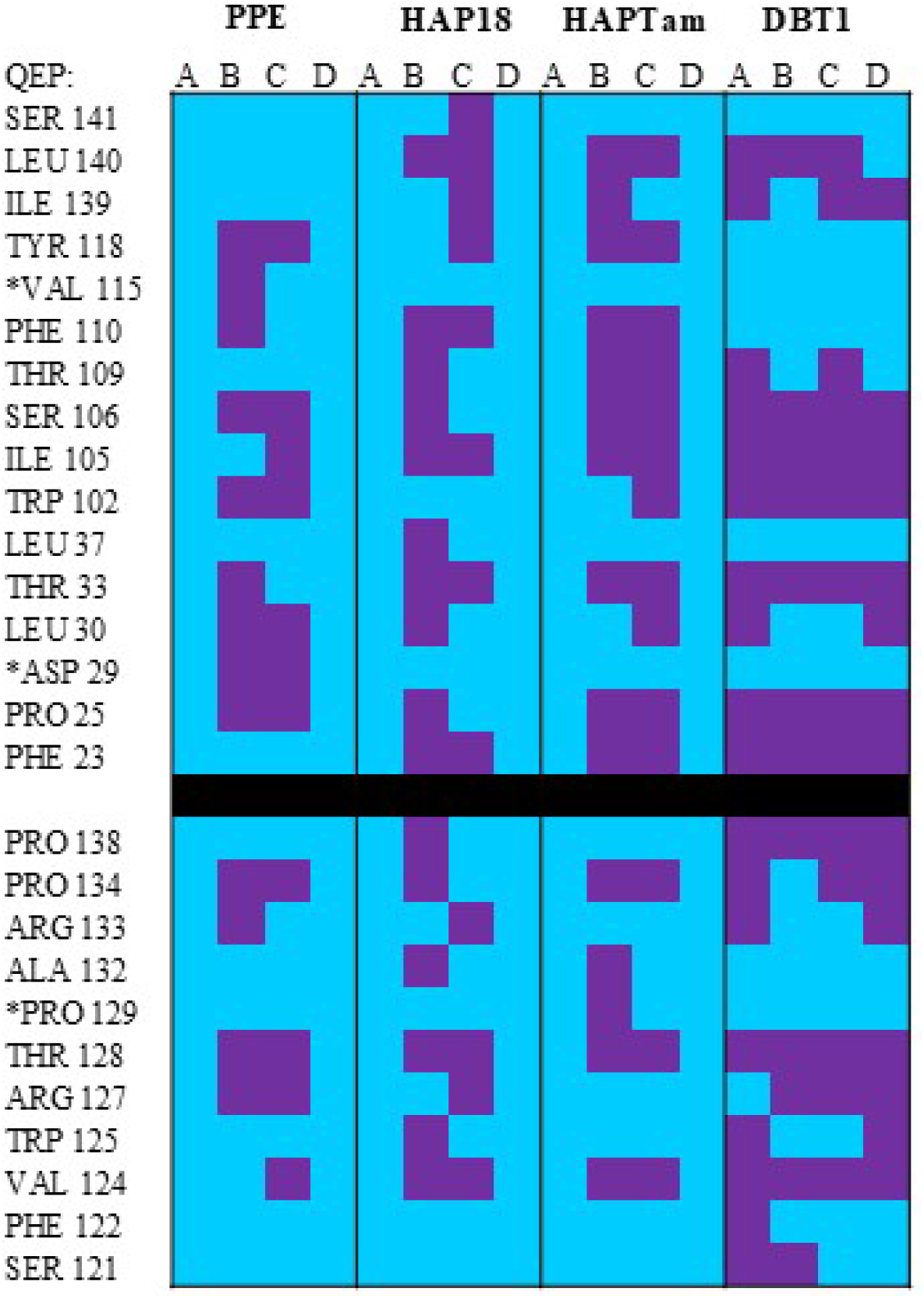
CAM interacting residues: Cp150. Residues with a 0.5 Angstrom VDW overlap with bound CAMs in Cp150 HBc structures. Purple = interacting, cyan = not interacting. (**Supplemental Table 1**)

**Supplemental Table 3:**
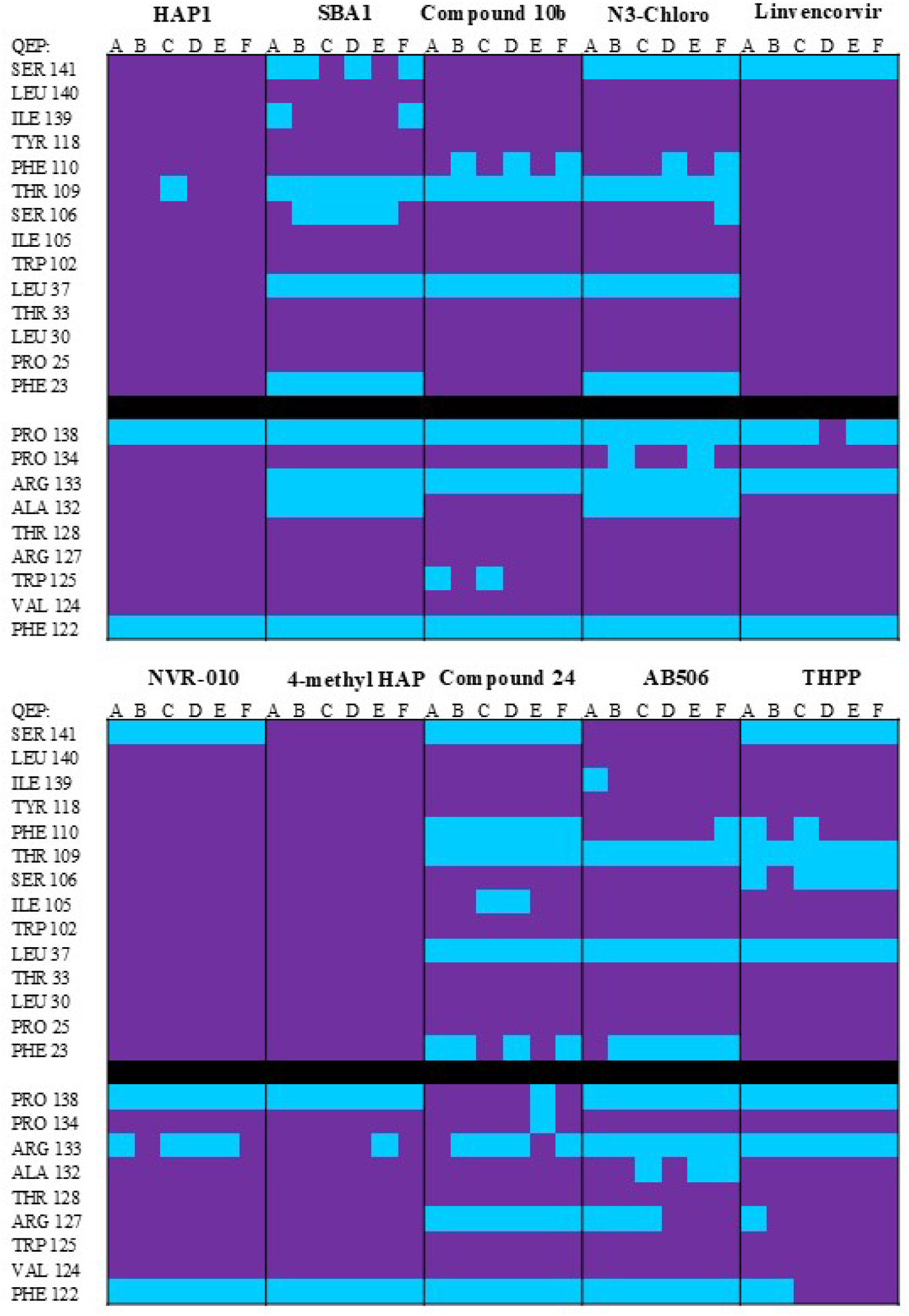
CAM interacting residues: Cp149-Y132A. Residues with a 0.5 Angstrom VDW overlap with bound CAMs in Cp149-Y132A HBc structures. Purple = interacting, cyan = not interacting. (**Supplemental Table 1**)

**Supplemental Table 4:**
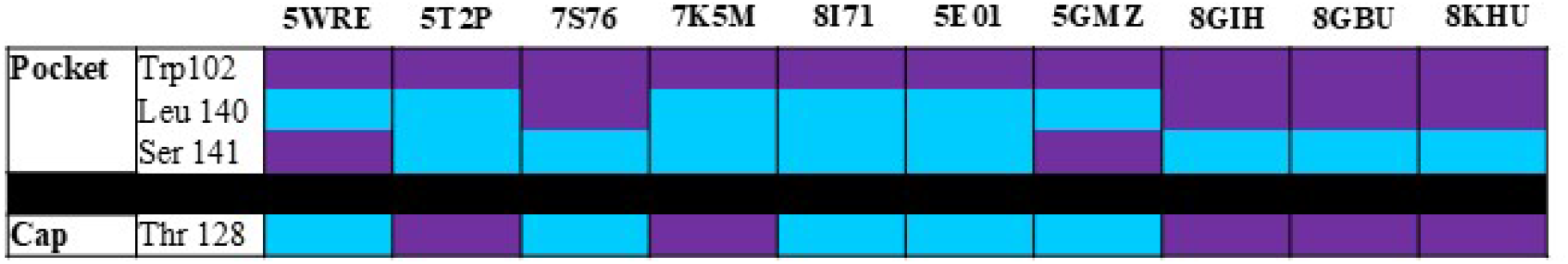
Residues that form hydrogen bonds with CAMs: Cp149-Y132A. Residues that form hydrogen bonds with bound CAMs in Cp149-Y132A HBc structures. For each structure, the CAM formed hydrogen bonds with the specified residues in all 6 available quasi-equivalent CAM sites. Purple = interacting, cyan = not interacting.(**Supplemental Table**

**Supplemental Table 5:**
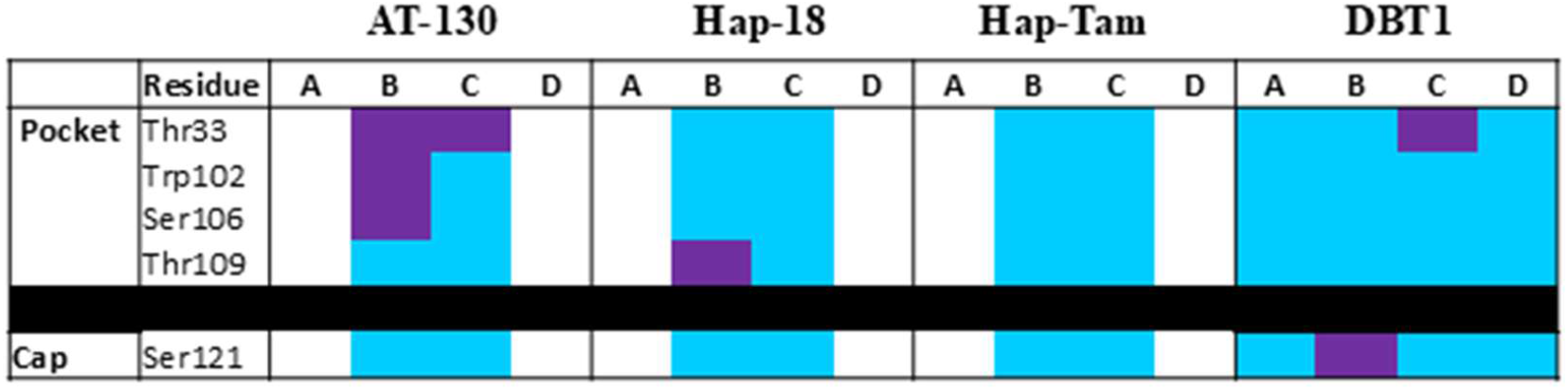
Residues that form hydrogen bonds with CAMs: Cp150. Residues that form hydrogen bonds with bound CAMs in Cp150 HBV-Cp structures. Purple = interacting, cyan = not interacting. Blank columns represent quasi-equivalent sites that are not CAM bound. (**Supplemental Table 1**)

